# pymfinder: a tool for the motif analysis of binary and quantitative complex networks

**DOI:** 10.1101/364703

**Authors:** Bernat Bramon Mora, Alyssa R. Cirtwill, Daniel B. Stouffer

**Affiliations:** Centre for Integrative Ecology, School of Biological Sciences, University of Canterbury, Christchurch, New Zealand; Department of Physics, Chemistry, and Biology, Linköping University, Linköping, Sweden.

## Abstract

We developed *pymfinder*, a new software to analyze multiple aspects of the so-called network motifs—distinct *n*-node patterns of interaction—for any directed, undirected, unipartite or bipartite network. Unlike existing software for the study of network motifs, *pymfinder* allows the computation of node- and link-specific motif profiles as well as the analysis of weighted motifs. Beyond the overall characterization of networks, the tools presented in this work therefore allow for the comparison of the “roles” of either nodes or links of a network. Examples include the study of the roles of different species and/or their trophic/mutualistic interactions in ecological networks or the roles of specific proteins and/or their activation/inhibition relationships in protein-protein interaction networks. Here, we show how to apply the main tools from *pymfinder* using a predator-prey interaction network from a marine food web. *pymfinder* is open source software that can be freely and anonymously downloaded from https://github.com/stoufferlab/pymfinder, distributed under the MIT License (2018).

## Introduction

The use of network theory has proven insightful in multiple fields, from the study of the spread of disease epidemics (1) to the characterization of neuronal networks (2). In ecology, this approach has been crucial to understanding the ways different species interact with each other, and the network perspective has justly become a central topic in community ecology (3). Over recent years, multiple methods for studying the topology of ecological networks have been successfully developed. Examples include models to generate realistic ecological communities (4) or tools for studying different network metrics such as compartmentalization (5), nestedness (6) or intervality (7). Following these advances, one of the most versatile ways to understand the structure of complex ecological networks is via the so-called network motifs—i.e. the analysis of small subgraphs representing the distinct patterns of interaction involving any set of *n* species. These subgraphs have been referred to as the ‘building blocks’ of complex networks (8).

The study of network motifs has been applied to multiple ecological systems over the recent years, including those composed of trophic (9) and mutualistic interactions (10). Non-ecological examples include in protein-protein interaction networks (11) and transcriptional regulation networks (12). There are typically two main approaches that are taken involving network motifs. First, counting the number of appearances of any given *n*-node pattern of interactions provides an overall perspective of the structure of a network. This has been done in different ecological studies, including the characterization of food webs (13), plant-pollination (14) and host-parasitoid networks (15). Second, other ecological studies have focused on the role of different species (16) and interactions (17), defining their position within the network based on which network motifs they form a part of. Following this work on network motifs, multiple tools for the counting of network motifs have been developed over the last decades (18, 19, 20). Most of the methodological work has focused on providing tools to efficiently quantify the overall structure of directed and undirected unipartite networks—i.e. graphs consisting of one set of interacting nodes. Unfortunately, to our knowledge, we are still lacking general-purpose software to also analyze bipartite networks—i.e. graphs consisting of two interacting sets of non-overlapping nodes—as well as to quantify the node-and link-specific motif profiles in both unipartite and bipartite networks. In addition, there is no tool to date that allows the user to include information regarding the interaction strengths of a network within the analysis of motifs. In response, we present *pymfinder*, software for motif analysis of network structure plus of the nodes and links of any type of network—i.e. directed/undirected, bipartite/unipartite, and weighted/binary networks.

*pymfinder* is an open-source and versatile tool for the study of network motifs and the result of long-standing research involving the study of ecological networks. For example, *pymfinder* was used to shed light on the ecological mechanisms underlying food-web structure (9), which, together with Bascompte 2005 (21) and Camacho et. al. (22), was one of the first studies to put network motifs into a purely ecological context. Building on these foundational studies, network motifs and *pymfinder* were shown to provide a useful way to characterize species’ roles, showing them to be evolutionary conserved across communities (16). Similarly, the roles of links involving parasite species were characterized through the study of network motifs, generating an understanding of how different types of feeding links are distributed within a food web (17). The same software has also been used to study bipartite networks. For instance, a study on host-parasitoid networks showed how species’ roles seem to be conserved over spatial scales as well as consistent over time (15). Perhaps more importantly, the software presented here has also been a central piece of very recent research. For example, the tools in *pymfinder* were used to relate species’ roles to multiple ecological traits in five marine food webs, showing that feeding environment is particularly strongly related to such roles (23). Likewise, the variability of species’ roles in plant-pollinator communities in the Arctic has recently shown to be related to the variability in community composition (24). Finally, the description of species’ roles has also been key to comparing entire networks by means of aligning species to each other, resulting in the identification of common backbones shared across food webs form different ecosystems (25). Overall, the tools included in *pymfinder* are and have been instrumental to the development of a diverse set of projects over the years, and we believe that they have the potential to be valuable for many others. This article describes the main structure of *pymfinder* and showcases some of its principal applications using a detailed ecological dataset as the backdrop.

## Design and implementation

### General description

*pymfinder* is a Python library that combines Python methods for network-motif analysis. Some of the engine underneath is a modified version of *mfinder*—a software tool for network-motif detection developed by Kashtan et. al. (8, 18). Originally, *mfinder* was written in C and made available solely as an executable, and we use it within *pymfinder* for its underlying efficiency. The *mfinder* code has been both included and modified here with the explicit consent of Nadav Kashtan, the author of mfinder 1.2.

As input, *pymfinder* accepts any type of network. That is, the analyses can be performed for both unipartite and bipartite networks. The format in which the networks are passed to the different functions of the package is either as text files, Python arrays or pymfinder-objects. Text files must describe the set of links comprising the networks, where each link appears as a separate line in the files. For example, a given line “*A B w*” would describe a single link *A* → *B* between nodes *A* and *B*, where *w* represents the strength or weight associated to such link (see Appendix). Similarly, Python arrays need to represent the list of interactions forming the networks. Notice that the direction of the links is important. Therefore, in bipartite networks, nodes of each group need to consistently be placed on the same side of the interactions—e.g. in a plant-pollinator networks the direction of the interactions in the input must all go from a plant to a pollinator (or vice-versa). Importantly, undirected networks can also be analyzed by *pymfinder*; however, any links between two nodes *A* and *B* in such networks need to be characterized by the two parallel links *A B* and *B A*. The output of *pymfinder*, is a high-level data type (‘class’) that contains different descriptors of the motif composition of the network under study (see Appendix).

### Structure of the package

At their core, all of the analyses performed by *pymfinder* are based around the identification of all the different *n*-node patterns of interaction found within a given network. To do this, *pymfinder* will always start by enumerating the unique motifs/subgraphs that make up the overall structure of the network under study. This analysis can be performed for multiple motif sizes. This is especially important for bipartite networks, where three-node motifs are minimally informative and one needs to explore bigger motifs (15). Notice, however, that increasing the number of nodes can be computationally challenging for unipartite networks since the number of unique motifs quickly increases with their size—i.e. there are 13 unique three-node motifs, 199 unique four-node motifs and 9364 unique five-node motifs.

For the sake of simplicity, we will focus most of the description of the methods presented here on the analysis of three-node network motifs. For any given network, this analysis is a three-step process. First, *pymfinder* loops through all the rows *i* of the adjacency matrix *A* associated with the network. For each non-zero element *a_ij_* found in row *A_i_*, it then searches for any connected element *a_jk_* = 1, *a_kj_* = 1, *a_ik_* = 1, and/or *a_ki_* = 1, revealing the existence of any motif comprised of the nodes *i*, *j*, and *k*. If *i*, *j* and *k* define a motif and this motif has not already been identified, the corresponding motif and the position of each node within the motif is recorded.

Based on this initial motif enumeration, *pymfinder* can perform three subsequent analyses: (i) the analysis of the overall network structure, (ii) the nodes and links’ participation in the different motifs, and (iii) the nodes and links’ role in each of the motifs.

#### Motif structure

The most basic application of *pymfinder* is the analysis of the overall motif structure of a given network. In particular, such analysis generates a description of the distribution of distinct *n*-node patterns of interaction found within the network (up to 8-node motifs). The application also includes the possibility of estimating the null motif composition expected for such network (see Appendix). To generate this null composition, *pymfinder* uses an MCMC algorithm to perform a randomization of the network while preserving the in- and out-degree of the nodes and each node’s number of single and double links (26, 27). Comparing the observed motif frequency to the random expectation, the application can be used to determine which interaction patterns are over- or under-represented relative to this null model (9). To do so, *pymfinder* calculates the mean and standard deviation of the null expectation as well as the z-scores for its comparison with the actual observations.

An additional feature of *pymfinder* is the possibility of incorporating information regarding the link strength into the analysis of the motif structure. This is notable in particular since there is no software available to explore the way the interaction strengths are distributed within networks across motifs. To do so, *pymfinder* will account for each motif within a given weighted network as a function of the strength of the links forming them (Fig. 1). Note that the algorithm allows the user to choose how the weight of a motif is defined. Specifically, given a motif formed by the set of links with strengths {*l*} = {*l*_1_,*l*_2_,…,*I_L_*}, *pymfinder* will calculate the weight of such motif as *f*({*l*}), where *f* is the function defined by the user. By default, *pymfinder* uses the arithmetic mean as the function *f*. Similar to unweighted networks, analysis of the motif structure of a weighted network returns the average and standard deviation of the weight of each motif, as well as the median and the first and third quartiles.

**Figure 1:**
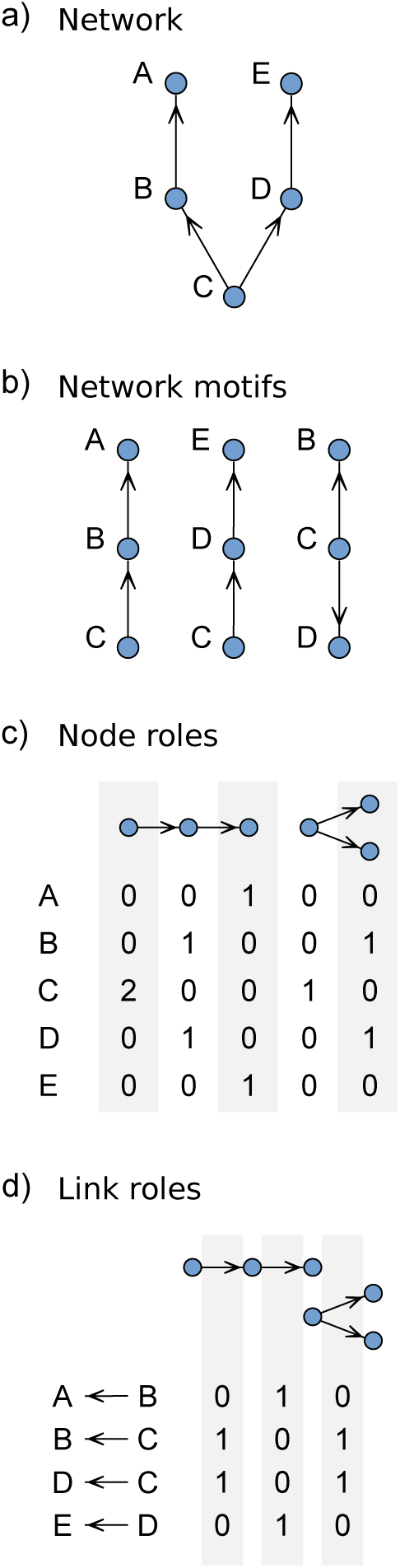
Main components of network-motif analysis. (a) A simple network that could represent a simple ecological community—where nodes would characterize species and the arrows would indicate the interactions between them—-or a protein-protein interaction network—where nodes would represent different proteins and the arrows indicate either activation or inhibition. (b) All three-node motifs found in the network from (a); from this classification, we can compute the overall network structure and the number appearances of every node in each motif. (c) The characterization of every node’s motif-role profile. This characterization is based on the number of appearances of every node in each of the unique motif node-positions. (d) The characterization of every link’s motif-role profile, which is based on the number of appearances of every link in each of the unique motif link-positions. Notice that we excluded any motif or role that was not represented in the network.

#### Motif participation

The study of network motifs can also be used as a way to classify nodes based on which patterns of interactions they are part of. For any given network, this application determines the frequency of appearance of every node across each of the different motifs (Fig. 1b), defining their participation across these distinct patterns of interactions. This a useful perspective for motif analysis because it provides a node-based description of the networks that can be used to understand the nature of specific nodes (e.g. different species in ecological networks or different proteins in protein-protein interaction networks) as well as decomposes the overall structure of the network at a finer resolution (21). Similarly, the same analysis can also be performed for the links forming the network. That is, *pymfinder* can quantify the frequency with which every link forms part of each distinct motif. As for the analysis of the overall structure of the networks, the motif participation of both nodes and links can also be calculated for any given motif size up to 8 nodes for weighted and unweighted networks. Again, *pymfinder* will account for each motif within a given weighted network as a function of the strength of the links forming it (Fig. 1), and the algorithm allows the user to choose this function just as described above for motif structure.

#### Motif-role profiles

Within any given motif, nodes can play multiple roles. For example, in the two-node motif *A* → *B*, there are two distinct positions *A* and *B*, which define two different roles—e.g. a predator and a prey in a food web. In contrast, for the two-node motif *A* ↔ *B*, *A* and *B* occupy indistinguishable positions; therefore, there is a single distinct role. The same idea can be extended to all *n*-node motifs. For example, there are 30 distinct node positions and 24 distinct link positions across the 13 unique three-nodes motifs. These distinct positions within the different motifs are important because the number of times that a node appears in each of them can be used as a way to define its structural role in a community (16). That is, we can characterize a node’s structural role based on the number of times that it occupies each distinct position of the *n*-node motifs. *pymfinder* provides a way to determine such *n*-node motif-role profiles for both the nodes (Fig. 1c) and the links (Fig. 1d) of a given network. Notice, however, that this function can only be run for two- and three-nodes motifs in unipartite networks, and two- to six-nodes motifs in bipartite networks.

The analysis of node and link motif-role profiles can also incorporate information regarding the strengths of interactions between nodes. As before, consider a motif *m* formed by the set of nodes {*i*} and the set of links with strengths {*l*}. For any node *j* in {*i*}, *pymfinder* calculates the contribution *C_jm_* of motif *m* to any of the positions of *j*’s motif-role profile as:

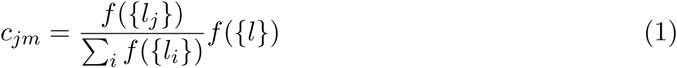

where {*l_i_*} is the set of strengths of all links in *m* involving node *i*, and *f* is a function defined by the user. By default, *pymfinder* again uses the arithmetic mean as *f* for weighted motif-role profiles. Notice that the contribution *c_im_* = 1 when ignoring the weights, or *f* is the arithmetic mean and all weights are equal to the motif size. When analyzing the motif-role profile of a link *k* forming such motif, the contribution *c_km_* is assumed to be exactly equal to its link strength *l_k_*.

### Basic tests

To ensure the reliable functioning of *pymfinder*, we included a set of basic tests in the package. All these basic tests are based around the idea of analyzing the structure of artificial networks containing only a single motif of each type for a given motif size—up to five-node motifs for bipartite networks and three-node motifs for unipartite networks. In addition, those networks are also set up so that any given node or link is only involved in a single motif and role. Using these single-motif networks, we tested the functions of *pymfinder* by ensuring that the analysis of such artificial networks does not result in the misrepresentation of any motif, node, link or role.

## Results/Discussion

The tools provided by *pymfinder* can be used in a large variety of systems and do not depend on the nature or providence of the networks. To illustrate the capabilities and potential of the software, we outline the study of a food web from a marine ecosystem as a representative study system (28). This specific network describes the predator-prey interactions between approximately 250 of the species found across an extensive area of the Caribbean Sea.

We first analyzed the overall three-species motif structure of the network and compared it to the random expectation (Fig. 2b). For this example, we used the z-score values to draw this comparison, which assume normality of the motif distribution. Notice, however, that *pymfinder* also returns the mean number of motif counts in the randomized networks, which allows for other types of statistical analyses. We found that the observed motif distribution is generally significantly different from the random expectation, showing either over- or under-representation relative to the results of the null model used here. This is evidence of a non-random organization of ecological communities (21, 29), which speaks to the eco-evolutionary mechanisms shaping the ways in which different species interact with each other. We then studied the distribution of link weights across motifs to test whether or not different motifs are generally made of different interaction strengths. For this particular example, we log-transformed the link weights to be approximately normally distributed as well as scaled them so that the weakest and strongest links had a weight of zero and one, respectively. In general, we found that interaction strengths are distributed in a similar manner across the different motifs of the network under study (Fig. 2c). Notice that these results are subject to the logarithmic transformation applied to the weight data, which is generally very skewed (28).

**Figure 2:**
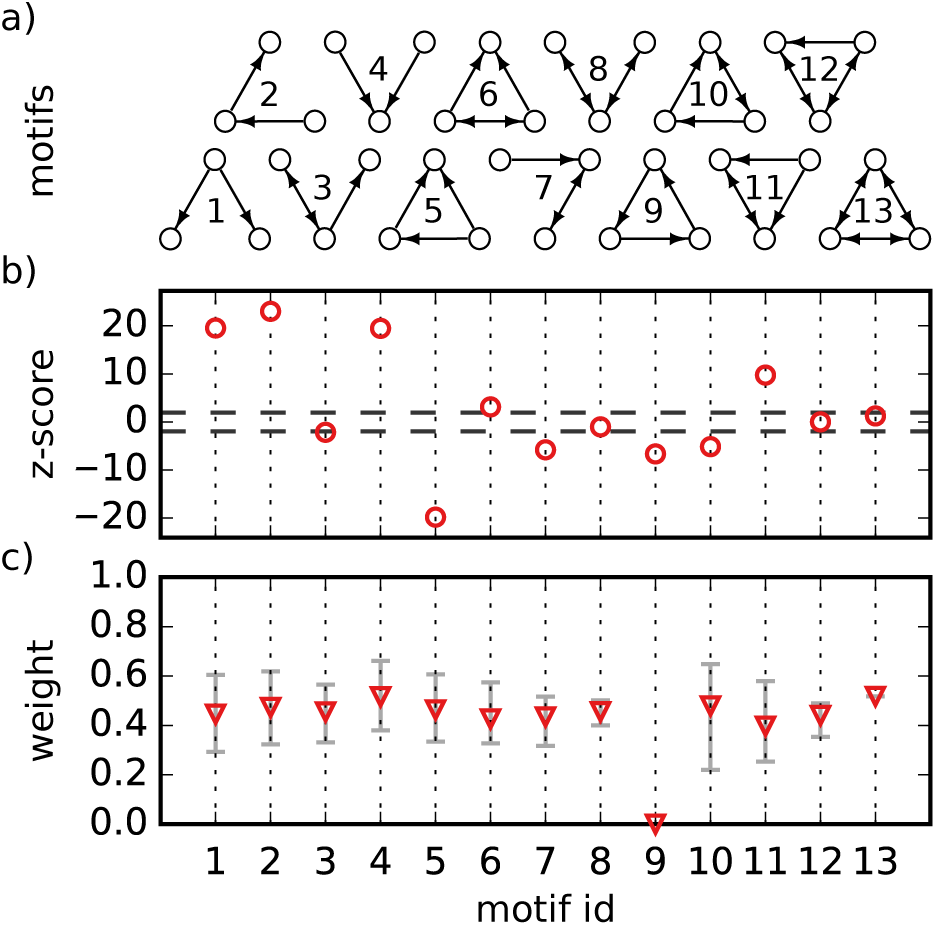
Analysis of the overall motif structure of the marine food web under study. The first panel (a) shows all the possible three-species motif structures. In this case, any arrow indicates the direction of energy flow from a prey to its predator. The second panel (b) presents the z-scores obtained from comparison between the empirical motif frequency and the random expectation. The dotted lines indicate the thresholds for significant over- and under-representation (*z* = 1.97 and *z* = –1.97, respectively). The third panel (c) shows the median weight found for each motif. The error bars represent the first and third quartiles. Note that the motif id given on the x-axis corresponds to the indexing in (a), and that the interaction strengths have been transformed to approximately be normally distributed and strictly positive.

Following the analysis of the overall motif structure, we examined the motif participation of the different nodes and links that make up this food web. We found that some nodes (e.g., sea cucumbers and algae) share almost identical motif-participation profiles while others (e.g., filefish and sea cucumbers) have very distinct profiles (Fig. 3a). This shows how motifs can be a valuable and insightful way to classify and compare the species across communities. Perhaps more importantly, we observed how the information regarding the interaction strengths forming the motifs changed those motif-participation profiles (Fig. 3b). Therefore, adding interaction strengths allowed us to distinguish between the roles of species with similar unweighted profiles. This is important because it suggests that, from a node-specific perspective, interaction strengths are not equally distributed across motifs. The uneven distribution of interaction strengths has important implications for the relationship between network structure and species–interaction strengths and the stability of food webs(30, 31). We also looked at the motif-participation profiles of the links (Fig. 3c). We found that those profiles could also be an indicator of the observed differences on the way interaction strengths are distributed across motifs, as suggested by previous work (17).

**Figure 3:**
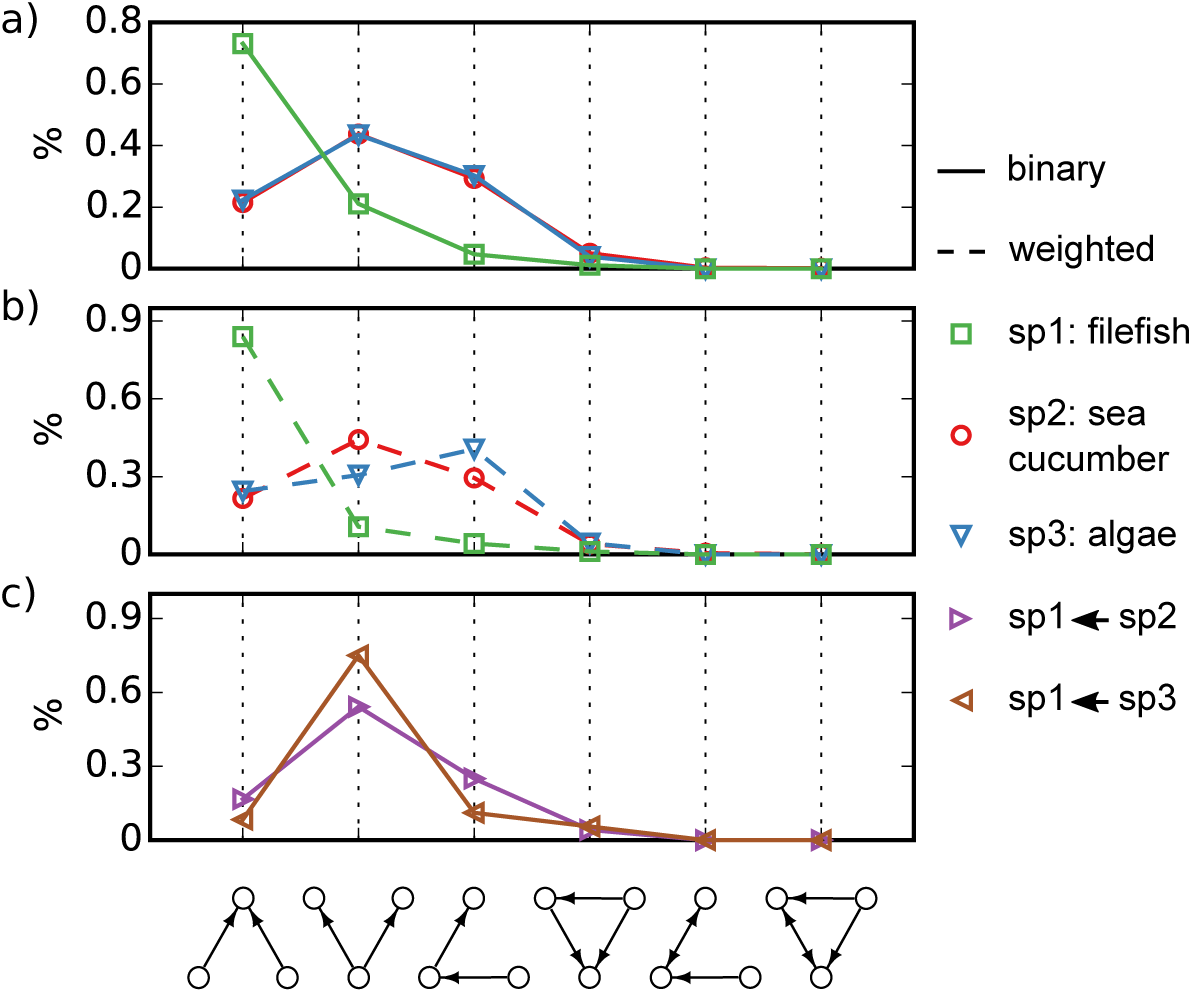
Analysis of the species’ motif participation in the marine food web under study. The first panel (a) shows the motif-participation profiles of three representative species from the web; here, every point describes the proportion of times that these species are found in any of the possible motifs. For simplicity, we excluded the seven motifs in which these species never appear. The second panel (b) presents the motif-role profiles for the same three species when adding information regarding the interaction strengths. In this case, every point represents the relative weight associated with the motifs in which each species participates. The third panel (c) shows the motif-participation profiles for the links involving the same three species.

Finally, we studied the motif-role profiles of the species of the marine network. This analysis is similar to the motif participation analysis of nodes and links; however, it provides a finer resolution to the role that different species or links might play in the community. Using the proportion of times that the different species are in each of the 30 unique positions of the three-species network motifs, we performed an analysis of multivariate homogeneity of group dispersions to compare the roles of the species in the network (32). To do this, we first calculated the euclidean distance between the roles of every pair of species in the network, generating a dissimilarity matrix of all species. We then performed a basic clustering analysis of the species-role dissimilarity matrix to find the most distinct groups of roles (Fig. 4). Finally, we used the function *betadisper* from the R package *vegan* (33) to perform the Principal Coordinates Analysis (PCoA) of the data. We found four characteristic groups of species presenting very distinct motif-role profiles. Notice that the same analysis can also be done for the motif-role profile of every link in the network. This is useful because it shows the diversity of structural roles in this community and underlines how those profiles could be used to compare species, links or networks within and across ecosystems, environments and biomes (34, 25).

**Figure 4:**
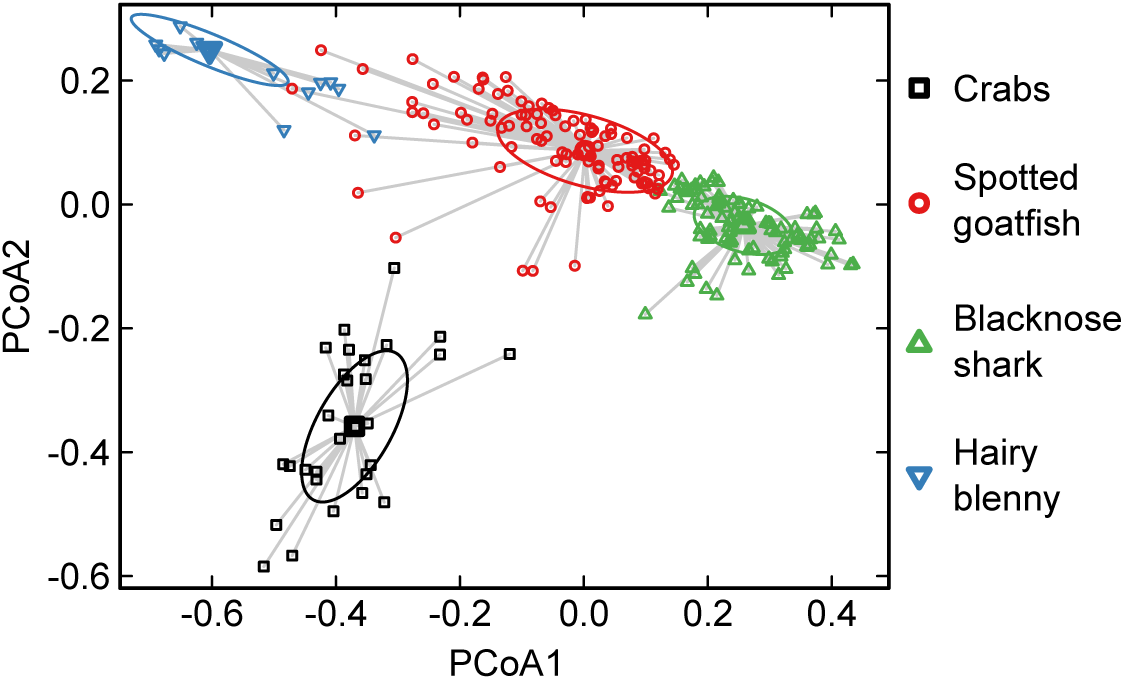
Principal coordinate analysis of the dissimilarity matrix containing the pairwise distances between all nodes’ motif-role profiles in the marine food web under study. Every point represents a different species and each color corresponds to a group characterizing a distinct role. The species in the legend are those corresponding to the medoids of each group. The ellipses are the one-standard-deviation ellipses about the group medians.

## Availability and Future Directions

*pymfinder* is open source software that can be freely and anonymously downloaded from https://github.com/stoufferlab/pymfinder. The documentation of the package is attached as supplementary material and the data used to test the software can be found within the github repository. *pymfinder* has been tested to run on any platform that supports Python. *pymfinder* will require you to have the Python modules Numpy and Setuptools installed in your machine. Data used to present the software has been previously published by Bascompte et. al. (28). We are currently working on additional software that uses the weighted motif-role profiles of nodes to efficiently align bipartite networks multiple times.

## References

[1] Mark EJ Newman. Spread of epidemic disease on networks. Physical review E, 66(1):016128, 2002.

[2] Olaf Sporns. Network analysis, complexity, and brain function. Complexity, 8(1):56–60, 2002.

[3] Jordi Bascompte and Pedro Jordano. Mutualistic networks. Princeton University Press, 2013.

[4] Richard J Williams and Neo D Martinez. Simple rules yield complex food webs. Nature, 404(6774):180–183, 2000.

[5] Roger Guimera, Marta Sales-Pardo, and Luís A Nunes Amaral. Module identification in bipartite and directed networks. Physical Review E, 76(3):036102, 2007.

[6] Jordi Bascompte, Pedro Jordano, Carlos J Melián, and Jens M Olesen. The nested assembly of plant–animal mutualistic networks. Proceedings of the National Academy of Sciences, 100(16):9383–9387, 2003.

[7] Daniel B Stouffer, Juan Camacho, and Luís A Nunes Amaral. A robust measure of food web intervality. Proceedings of the National Academy of Sciences, 103(50):19015–19020, 2006.

[8] Ron Milo, Shai Shen-Orr, Shalev Itzkovitz, Nadav Kashtan, Dmitri Chklovskii, and Uri Alon. Network motifs: simple building blocks of complex networks. Science, 298(5594):824–827, 2002.

[9] Daniel B Stouffer, Juan Camacho, Wenxin Jiang, and Luís A Nunes Amaral. Evidence for the existence of a robust pattern of prey selection in food webs. Proc R Soc Biol Sci, 274(1621):1931–1940, 2007.

[10] Carsten F Dormann, Jochen Fründ, Nico Blüthgen, and Bernd Gruber. Indices, graphs and null models: analyzing bipartite ecological networks. The Open Ecology Journal, 2(1), 2009.

[11] Esti Yeger-Lotem, Shmuel Sattath, Nadav Kashtan, Shalev Itzkovitz, Ron Milo, Ron Y Pinter, Uri Alon, and Hanah Margalit. Network motifs in integrated cellular networks of transcription–regulation and protein–protein interaction. Proceedings of the National Academy of Sciences of the United States of America, 101(16):5934–5939, 2004.

[12] Shai S Shen-Orr, Ron Milo, Shmoolik Mangan, and Uri Alon. Network motifs in the transcriptional regulation network of escherichia coli. Nature genetics, 31(1):64, 2002.

[13] Janis Klaise and Samuel Johnson. The origin of motif families in food webs. Scientific Reports, 7(1):16197, 2017.

[14] María C Rodriíguez-Rodríguez, Pedro Jordano, and Alfredo Valido. Functional consequences of plant-animal interactions along the mutualism-antagonism gradient. Ecology, 98(5):1266–1276, 2017.

[15] Nick J Baker, Riikka Kaartinen, Tomas Roslin, and Daniel B Stouffer. Species’ roles in food webs show fidelity across a highly variable oak forest. Ecography, 38(2):130–139, 2015.

[16] Daniel B Stouffer, Marta Sales-Pardo, M Irmak Sirer, and Jordi Bascompte. Evolutionary conservation of species’ roles in food webs. Science, 335(6075):1489–1492, 2012.

[17] Alyssa R Cirtwill and Daniel B Stouffer. Concomitant predation on parasites is highly variable but constrains the ways in which parasites contribute to food web structure. Journal of Animal Ecology, 84(3):734–744, 2015.

[18] Nadav Kashtan, Shalev Itzkovitz, Ron Milo, and Uri Alon. Efficient sampling algorithm for estimating subgraph concentrations and detecting network motifs. Bioinformatics, 20(11):1746–1758, 2004.

[19] Sebastian Wernicke and Florian Rasche. Fanmod: a tool for fast network motif detection. Bioinformatics, 22(9):1152–1153, 2006.

[20] Gabor Csardi and Tamas Nepusz. The igraph software package for complex network research. InterJournal, Complex Systems, 1695(5):1–9, 2006.

[21] Jordi Bascompte and Carlos J Melian. Simple trophic modules for complex food webs. Ecology, 86(11):2868–2873, 2005.

[22] Juan Camacho, Daniel B Stouffer, and Luís A Nunes Amaral. Quantitative analysis of the local structure of food webs. Journal of Theoretical Biology, 246(2):260–268, 2007.

[23] Alyssa R Cirtwill and Anna Eklof. Feeding environment and other traits shape species’ roles in marine food webs. Ecology letters, 21(6):875–884, 2018.

[24] Alyssa R Cirtwill, Tomas Roslin, Claus Rasmussen, Jens Mogens Olesen, and Daniel B Stouffer. Between-year changes in community composition shape species’ roles in an arctic plant–pollinator network. Oikos, 2018.

[25] Bernat Bramon Mora, Dominique Gravel, Gilarranz J. Luis, Poisot Thimothee, and Daniel B. Stouffer. Identifying a common backbone of interactions underlying food webs from different ecosystems. Nature Communications, 2018. (accepted).

[26] Ron Milo, Nadav Kashtan, Shalev Itzkovitz, Mark EJ Newman, and Uri Alon. On the uniform generation of random graphs with prescribed degree sequences. arXiv preprint cond-mat/0312028, 2003.

[27] James G Sanderson, Michael P Moulton, and Ralph G Selfridge. Null matrices and the analysis of species co-occurrences. Oecologia, 116(1-2):275–283, 1998.

[28] Jordi Bascompte, Carlos J Melián, and Enric Sala. Interaction strength combinations and the overfishing of a marine food web. Proceedings of the National Academy of Sciences of the United States of America, 102(15):5443–5447, 2005.

[29] Jonathan J Borrelli. Selection against instability: stable subgraphs are most frequent in empirical food webs. Oikos, 124(12):1583–1588, 2015.

[30] Anje-Margriet Neutel, Johan AP Heesterbeek, and Peter C de Ruiter. Stability in real food webs: weak links in long loops. Science, 296(5570):1120–1123, 2002.

[31] Mark Emmerson and Jon M Yearsley. Weak interactions, omnivory and emergent food-web properties. P Roy Soc Lond B Bio, 271(1537):397–405, 2004.

[32] Marti J Anderson. Distance-based tests for homogeneity of multivariate dispersions. Biometrics, 62(1):245–253, 2006.

[33] Jari Oksanen, F. Guillaume Blanchet, Michael Friendly, Roeland Kindt, Pierre Legendre, Dan McGlinn, Peter R. Minchin, R. B. O’Hara, Gavin L. Simpson, Peter Solymos, M. Henry H. Stevens, Eduard Szoecs, and Helene Wagner. vegan: Community Ecology Package, 2017. R package version 2.4–4.

[34] Timothée Poisot, Elsa Canard, David Mouillot, Nicolas Mouquet, and Dominique Gravel. The dissimilarity of species interaction networks. Ecol Lett, 15(12):1353–1361, 2012.

